# Safety assessment of insecticidal proteins: A case study utilizing His-tagged protein to determine the safety of Mpp75Aa1.1, a new ETX_MTX2 protein that controls western corn rootworm

**DOI:** 10.1101/2022.03.28.486136

**Authors:** Cunxi Wang, Gregory J. Bean, Chun Ju Chen, Colton R. Kessenich, Jiexin Peng, Nicolo R. Visconti, Jason S. Milligan, Robert G. Moore, Jianguo Tan, Thomas C. Edrington, Bin Li, Kara S. Giddings, David Bowen, Jinhua Luo, Todd Ciche, William J. Moar

**Affiliations:** Bayer Crop Science, 700 Chesterfield Pkwy West, Chesterfield, MO 63017

**Keywords:** Mpp75Aa1.1, GM crops, β pore forming protein, His-tagged protein, safety assessment

## Abstract

The recently discovered insecticidal protein Mpp75Aa1.1 from *Brevibacillus laterosporus* is one member of the ETX_MTX family of beta-pore forming proteins (β-PFPs) expressed in genetically modified (GM) maize to control western corn rootworm (WCR; *Diabrotica virgifera virgifera* LeConte). In this paper, bioinformatic analysis establishes that although Mpp75Aa1.1 shares varying degrees of similarity to members of the ETX_MTX2 protein family, it is unlikely to contribute to any allergenic, toxic, or otherwise adverse biological effects. The safety of Mpp75Aa1.1 is further supported by a weight of evidence including evaluation of history of safe use (HOSU) of ETX_MTX2 proteins. Assessments using either purified Mpp75Aa1.1 protein or a poly-histidine-tagged (His-tagged) variant of the Mpp75Aa1.1 protein demonstrate that both forms of the protein are heat labile at temperatures at or above 55 °C, degraded by gastrointestinal proteases within 0.5 min, and have no adverse effects in acute mouse oral toxicity studies at a dose level of 1920 or 2120 mg/kg body weight. Results also indicate that the domain-based protein characterization should be considered as part of the weight of evidence for the safe consumption of food or feed. Furthermore, these results support the use of His-tagged proteins as suitable surrogates for assessing the safety of their non-tagged parent proteins.

## 1 INTRODUCTION

Insect pests in agricultural systems are a major cause of damage to crop production worldwide, with productivity losses estimated from 26 – 80% (1) and economic losses of 300 billion USD (2). Historically, pest control in industrialized countries has relied on the use of synthetic chemical pesticides (3). However, the increased insect resistance has stimulated significant research and development efforts to identify new biological pesticides. Microbial-derived pesticides provide an environmentally-friendly approach that has been used in crop protection for nearly a century, including in modern organic farming (4). Because of their efficacy towards the control of insect pests and strong human and environmental safety profile, insecticidal proteins isolated from naturally occurring bacteria (e.g., *Bacillus thuringiensis*, or *Bt*) have been safely utilized in genetically modified (GM) crops since the first commercialization in 1996 (5). GM crops expressing microbially-derived insecticidal protein(s) have been widely adopted and accepted as an important tool for farmers to control insect pests, accounting for 94% of planted soybean, 92% of planted maize and 96% of planted cotton in the United States in 2020 (4, 6–9).

The expanded use of GM crops expressing a limited number of insecticidal proteins has created intense selection pressure for the development of resistant insects (10). This has driven a critical need to discover and develop new insecticidal proteins to minimize the evolution of target insect resistance and expand the spectrum to include other insect pests. One such newly discovered insecticidal protein is Mpp75Aa1 derived from *Brevibacillus laterosporus,* demonstrating insecticidal activity against the important maize pest, western corn rootworm (WCR), *Diabrotica virgifera virgifera* LeConte (11). Transgenic maize plants expressing a mature form of Mpp75Aa1 lacking the native transit signal peptide, Mpp75Aa1.1, have demonstrated significant below-ground protection from WCR in field trials (11).

Proteins expressed in GM crops undergo a rigorous safety assessment prior to commercialization. The safety assessment of newly expressed proteins in GM crops follows an internationally recognized framework established by the Food and Agriculture Organization of the United Nations (FAO)/ the World Health Organization (WHO) Codex Alimentarius commission in 2009 (12). Under this framework, proteins are assessed using a tiered, weight of evidence based approach (5, 12–16) which has been extensively used to successfully determine the safety of GM-derived proteins for human and other vertebrate animal consumption. The first tier of this approach includes an evaluation of the history of safe use (HOSU) of the expressed protein, a characterization of its physicochemical and functional properties, an assessment of its susceptibility to digestive enzymes and its stability after heat treatment at temperatures representative of grain processing and cooking. In addition, another key step in Tier I is an *in-silico* assessment using bioinformatics. These bioinformatics analyses not only assess whether or not the introduced protein shows any similarity to known toxins or allergens, but also classify the structural and functional relationship of the newly expressed protein with previously identified proteins. The second tier of the weight of evidence evaluates the potential for mammalian toxicity *in vivo* (*13*).

Primary amino acid sequence and structural homology have identified Mpp75Aa1.1 as belonging to the *Clostridium epsilon* toxin ETX/*Bacillus* mosquitocidal toxin MTX2 (ETX_MTX2) protein family (Pfam:PF03318) (17, 18). The ETX_MTX2 protein family is one of β-pore forming protein (β−PFPs) families, characterized by a shared common mode of action (MOA) that utilizes an amphipathic hairpin loop to insert into cell membranes to form a β-barrel membrane-integrated pore. ETX_MTX2 β−PFPs are commonly produced by bacteria, fungi, plants, and some animals, and are present in many safely consumed foods such as spinach, cucumber, wheat, and fish (19).

In addition, the safe use of β−PFPs in GM crops for food and feed has also been established (15, 19). The Tpp35Ab1 protein (formerly classified as Cry35Ab1) is from the Toxin_10 family of β−PFPs (20). GM crops containing Tpp35Ab1 and its binary partner, Gpp34Ab1 (formerly classified as Cry34Ab), have been commercially planted for millions of acres in the U.S. corn belt since 2006 providing an extensive HOSU and exposure for human and other vertebrate animal consumption of β−PFPs (19). Notably, empirical mammalian safety data generated for both proteins, β−PFP safety (21). Safe consumption of ETX_MTX2 proteins was further evidenced by assessing Mpp51Aa2.834_16 (formerly classified as Cry51Aa2.834_16) using a weight of evidence approach (15).

Although structurally diverse from three-domain *Bt* Cry proteins, the MOA of ETX_MTX2 proteins follows the same general steps as the well-established *Bt* Cry proteins, including proteolytic activation, specific binding to brush border membranes followed by pore formation and insect death (22). While the β−PFPs are classified by their common structure/function, crystal structure analyses demonstrate that ETX_MTX2 proteins have structurally diversified receptor binding domains (23–25) that play an important role in their insecticidal activity. The results from the ETX_MTX2 protein Mpp51Aa2.834_16 confirmed that the functional specificity and selectivity against targeted insect pests within this protein family is conferred by specific residues within the receptor-binding domain (24, 26). Screening analysis has also suggested that the receptor-binding region of the Mpp75Aa1.1 protein, which is similarly diversified from other ETX_MTX2 proteins, is responsible for its insecticidal specificity (18).

To assess the safety of a newly expressed protein, hundreds of grams of purified test protein are required and must be produced using a multi-step purification process optimized for each new protein. For difficult to produce proteins, production of sufficient amounts of a protein at a suitable purity level can be limited towards conducting a thorough safety assessment (27). An affinity tag system can simplify the protein purification process, and in some instances improve the tractability of a protein of interest. For these reasons, affinity-tags such as the poly-histidine tag system (His tag) are often used to purify proteins for functional and biophysical studies. Typically the His tag is covalently attached to a protein at the N- or C-terminal amino acid and has a negligible effect on the native structure of the protein (28). Further, numerous studies have been conducted using His-tagged proteins, including but not limited to enzymatic activity assays (29–31), protein-protein interaction studies (32, 33), labeling (34, 35), vaccine expression and purification (36). While His-tagged proteins are used commonly for functional and structural characterization, the lack of data available regarding the impact of affinity tags on the safety profile of proteins has limited their use in safety assessments of newly expressed proteins in GM crops.

The present study focuses on the safety assessment of Mpp75Aa1.1. A bioinformatic analysis demonstrates that Mpp75Aa1.1 shares some sequence homology to known β-PFPs, however, the proteins differ in sequence and structure and that observation, together with the appropriate testing and assessments, indicates no tox or allergenicity concerns. In addition, the equivalence studies characterizing the safety of a protein with and without the presence of a His-tag suggest that affinity-tagged proteins are suitable surrogates for future protein safety studies.

## 2 MATERIALS AND METHODS

### 2.1 Expression and purification of Mpp75Aa1.1

The full-length precursor Mpp75Aa1 has a 23 amino acid N-terminal membrane transiting signal peptide (11). In this study, the mature form of Mpp75Aa1 without the N-terminal 23 amino acids was produced and accessed. To differentiate the full-length Mpp75Aa1 from its mature form, the mature form was assigned as Mpp75Aa1.1. The Mpp75Aa1.1 coding sequence with (-HHHHAHHH, thereafter referred to as Mpp75Aa1.1-His) or without the C-terminal His tag (thereafter referred as Mpp75Aa1.1) was cloned into a pET24a vector (Novagen, Madison, WI) and expressed in Rosetta 2 (*DE3*) *E. coli* (Invitrogen, Carlsbad, CA) cells. Cells were grown in auto-induction media (37) containing 50 mg/L Kan and 30 mg/L chloramphenicol at 37°C for 3-4 hr and then continually cultured at 20°C for ∼24 hr.

For small scale purification of Mpp75Aa1.1 and Mpp75Aa1.1-His (used for side-by-side comparison of functional activity, heat stability & digestibility), each protein was extracted from *E. coli* fermentation product using 10 mM sodium carbonate/bicarbonate, pH 10.8, 1 mM EDTA, 1 mM Benzamidine and 100 U/ml Benzonase (buffer A). The protein extracts were clarified by centrifugation, loaded onto a Q Sepharose Fast Flow column (GE healthcare, Piscataway, NJ) and eluted by step gradients starting with 50 mM NaCl and 25 mM increment in buffer A. The Mpp75Aa1.1-containing fractions were pooled, concentrated and buffer-exchanged into 10 mM sodium carbonate/bicarbonate pH 10 using a 12 to 14 kDa Spectra/Por® 2 Dialysis Membrane (Spectrum Laboratories, Inc.) followed by centrifugal concentration devices (Amicon® Ultra-15 Centrifugal Filter Unit, EMD Millipore Corporation, Billerica, MA). Both Mpp75Aa1.1 and Mpp75Aa1.1-His were stored at -80°C in a 10 mM sodium carbonate/bicarbonate, pH 10 buffer before use.

For large scale purification of Mpp75Aa1.1 (used for the acute toxicity study), inclusion bodies were collected from *E. coli* fermentation by centrifugation after cell disruption and washed extensively in a 3-step process, first with 50 mM Tris, 150 mM NaCl, (pH 8.0), then 50 mM sodium carbonate/bicarbonate, 2% Triton-X 100, 1 M NaCl, (pH 10.8), and lastly 10 mM sodium carbonate/bicarbonate, 5 mM DTT, (pH 10.8). Washed inclusion bodies were solubilized in 10 mM sodium carbonate/bicarbonate, (pH 10.8) and loaded on a Q Sepharose Fast Flow column (GE healthcare, Piscataway, NJ) and eluted by a single step gradient at 200 mM NaCl. Mpp75Aa1.1-containing fractions were pooled, concentrated, and buffer exchanged via a hollow fiber ultrafiltration apparatus (30 kDa Molecular Weight Cut-Off, GE healthcare, Piscataway, NJ) into a buffer containing 10 mM sodium carbonate/bicarbonate (pH 10) and then stored at -80°C before use. The resulting protein was 100% pure determined by SDSPAGE and the identity of this large batch Mpp75Aa1.1 was confirmed by determination of its N-terminal sequence and sequence coverage of the peptide mass fingerprint (38).

For large scale purification of Mpp75Aa1.1-His (used for the acute toxicity study), inclusion bodies were collected by centrifugation after cell disruption and washed extensively, then the protein was solubilized in a 100 mM sodium carbonate (pH 11) buffer, bound to HisSelect Ni resin (Sigma-Aldrich, St Louis, MO) in batch mode, poured into a gravity flow column, washed, and eluted with imidazole-containing buffer at pH 11. After elution, buffer exchange was by dialysis against a buffer of 15 mM CHES, pH 9.8, 25 mM NaCl, 0.25 mM CaCl_2_ for 12 hours with 3 time of buffer changes. The Mpp75Aa1.1-His fractions identified by SDS-PAGE were pooled and further processed using EndoTrap® HD (Hyglos GmbH, Germany) to reduce endotoxin contents (manufacturer’s instruction). The protein was stored in a buffer containing 15 mM CHES (pH 9.8), 25 mM NaCl, 0.25 mM CaCl_2_ at -80°C before use. The protein was 99% pure and the identity of this large batch Mpp75Aa1.1-His was confirmed by determination of its N-terminal sequence, C-terminal His tag and sequence coverage of the peptide mass fingerprint (38).

### 2.2 Assessment of the presence of Mpp75A in *B. laterosporus*

Two strains of *B. laterosporus* were isolated following methods previously described (11). The EG4227 strain was isolated from grain dust while the BOD strain was isolated from the powder of a commercially available probiotic capsule. Cell cultures for each strain and DNA extractions methods were previously reported (11, 18). The Mpp75A genes were cloned from the EG4227 and BOD strain, respectively and the two amino acid sequences, deduced from the gene sequence, were designated as Mpp75Ab1 and Mpp75Ab2 by the BPPRC nomenclature committee using methods previously described (11). Whole cell cultures grown for 40 hr in terrific broth (TB) from the 2 strains extracted were in 2X LDS loading buffer with 100 mM DTT at a 1:1 ratio and 15 μl was loaded per lane. Cell-free culture supernatants from 40 hr TB cultures were extracted in 2X LDS loading buffer with 100 mM DTT at a 1:1 ratio and 15 μl was loaded per lane. For western blot analysis, culture samples were subjected to SDS–PAGE using the Bio-Rad Criterion**^TM^** system on a 4-20% Tris/Glycine/SDS gel with the related Tris/Glycine/SDS running buffer at 250V for 30 minutes. The proteins were electro-transferred from the gel to 0.2 μM nitrocellulose membrane using Bio-Rad Turbo Blot**^TM^** device for 7 minutes. The membrane was probed with a primary anti-mMpp75Aa2 (mature form of Mpp75Aa2, (11)) polyclonal antibody diluted at 1 μg/ml in the 1X phosphate buffered saline with 0.1% Tween **^®^** 20 with PBST + 2% non-fat milk (NFMK) overnight (∼12 hours at 4°C). The membrane was washed 3 times with 1X PBST to remove primary antibody. The membrane was probe with the secondary goat anti-rabbit (antibody-horse radish peroxidase conjugate) at 1:200,000 dilution in 2% NFMK in PBST for 1 hour at 4°C. The immunoreactive bands were detected using Supersignal West Femto**^TM^** maximum sensitivity kit (Pierce^TM^).

### 2.3 Characterization of Mpp75Aa1.1

The methods used to characterize the Mpp75Aa1.1 have previously been reported (38, 39). The concentration of Mpp75Aa1.1-His was determined by amino acid analysis. The concentration of total protein in the Mpp75Aa1.1 sample was determined by A280 using the Mpp75Aa1.1-His protein as a standard. The purity and apparent molecular weight of each protein was determined by densitometric analysis of Coomassie stained SDS–PAGE gels. A Q-exactive mass spectrometry (MS) (Thermo Fisher) or Orbitrap Fusion (Thermo Fisher) was used to confirm protein sequence identity. For mass analysis, aliquots of the protein samples were separated by SDS -PAGE gels. The bands corresponding to Mpp75Aa1.1 or Mpp75Aa1.1-His were excised from the gel and digested by trypsin after destaining, reduction and alkylation procedures. The tryptic peptides were extracted, dried down and re-dissolved into 2% acetonitrile, 0.1% formic acid in water, and injected into the MS for analysis. The identified peptides were used to assemble a sequence coverage. For intact mass analysis, the intact mass was collected after injection of an aliquot of samples and verified against the theoretic mass.

### 2.4 Bioinformatic assessment of Mpp75Aa1.1 allergenicity and toxicity

Bioinformatic assessments and thresholds used to assess any potential for allergenicity and/or toxicity were derived from those described previously by Wang et al. (2015) and outlined by Codex Alimentarius (2009). The database used to represent all known proteins was a download of all proteins in GenBank release 235 (40). An updated toxin database (herein described as TOX_2020) was built using the query “(keyword:toxin OR annotation:(type:"tissue specificity" venom)) AND reviewed:yes” to search the Swiss-Prot database (https://www.uniprot.org/uniprot/) to isolate likely toxins based on sequence descriptions (41). A second step was applied where the collected sequences were counter-screened to remove unlikely toxins on the basis of descriptions such as “antitoxin” or “non-toxic”. The end result is that the TOX_2020 toxin database contains 7,728 sequences. The allergen database (herein described as AD_2020) utilized was the "COMprehensive Protein Allergen REsource" (COMPARE) database as generated and maintained by the Health and Environmental Sciences Institute (HESI) (42), and contains 2,248 sequences. Alignments were generated using FASTA v36.3.5d run with an *E*-score cutoff of 1, and a threshold of ⩽1e−5 (1 × 10^−5^) was used as an initial threshold for alignment significance. This is the same threshold as used previously (38, 39) and is recognized as being a conservative threshold for the identification of proteins that may be homologous (43).

### 2.5 Assessment of Mpp75Aa1.1 susceptibility to pepsin

The susceptibility of Mpp75Aa1.1 to degradation by pepsin was assessed following a standardized protocol (44). Briefly, Mpp75Aa1.1 or Mpp75Aa1.1-His was mixed with high purity pepsin (Sigma, St. Louis, MO) in 2 mg/ml NaCl, 10 mM HCl, pH ∼1.2 to a final protein-to-pepsin ratio of 1 μg total protein:10 U of pepsin. The reaction mixture tube was immediately placed in a 37 ± 2°C water bath. Samples were removed at 0.5, 2, 5, 10, 20, 30 and 60 min and were immediately quenched by the addition of sodium carbonate and 5X SDS-PAGE sample loading buffer (∼310 mM Tris-HCl, 25% (v/v) 2-mercaptoethanol, 10% (w/v) sodium dodecyl sulfate, 0.025% (w/v) bromophenol blue, 50% (v/v) glycerol, pH 6.8). Protein only and pepsin only experimental controls were also prepared and incubated for 60 min in a 37 ± 2°C water bath. All resulting samples were heated at 95-100 °C for 5-10 mins, frozen on dry ice, and stored in a -80 °C freezer prior to SDS-PAGE analysis.

The extent of test protein digestion was assessed by both Brilliant Blue G staining of SDS-PAGE gels. In each case, the limit of detection (LOD) of the test proteins was determined.

### 2.6 Heat lability of Mpp75Aa1.1

In a method development assay, Mpp75Aa1.1 and Mpp75Aa1.1-His treated at 95°C completely lost activity while protein incubated at 25°C remained fully functional. Therefore, in this study, three temperatures from 37, 55 and 75°C along with a control treatment on wet ice were selected to evaluate the response of Mpp75Aa1.1 and Mpp75Aa1.1-His to heating by comparing their functional activities against WCR. Test proteins were exposed to each temperature for 15 min followed by storage on wet ice. The control sample aliquots of Mpp75Aa1.1 and Mpp75Aa1.1-His were maintained on wet ice throughout the course of the heat treatment incubation period. Following heat treatment, all samples were tested in WCR diet overlay bioassays to assess the insecticidal activity.

Insecticidal activity of heat-treated Mpp75Aa1.1 and Mpp75Aa1.1-his samples were evaluated using WCR eggs provided by the Bayer lab at Waterman, IL. Eggs were incubated at target temperatures ranging from approximately 10°C to 27°C to obtain the desired hatch time. Newly-hatched WCR larvae (≤30 hours after the first observation of hatching) were used in all bioassays. Mpp75Aa1.1 samples were prepared by diluting samples with a buffer solution of 10 mM sodium carbonate and bicarbonate, pH 10.0. For comparison of Mpp75Aa1.1 and Mpp75Aa1.1-His functional activity, bioassays consisted of a series of 9 dilutions yielding a concentration series with a two-fold separation factor ranging from 5 to 1200 µg/ml. All other bioassays consisted of a series of seven dilutions yielding a concentration series with a two-fold separation factor ranging from 7.8 to 500 µg/ml for treatments on wet ice and at 37°C and from 94 to 6000 µg/ml for treatment at 55°C and 75°C. For each bioassay, the concentration series was expected to elicit a response from WCR larvae allowing for determination of the LC_50_ value. The bioassay included buffer control that contained a buffer of the same composition used to suspend Mpp75Aa1.1.

Twenty µl of Mpp75Aa1.1 solution or buffer control was overlaid onto the surface of WCR diet in 96 well plates (Falcon). WCR diet was prepared according to manufacturer’s guidelines for SCR diet (Bio-Serv, Frenchtown, NJ) with supplements of formalin at 0.06% (v/v), 10% KOH (v/v) to adjust diet pH to 9, and lyophilized conventional corn root tissue at 0.62% (w/v). Two hundred μl of molten diet was pipetted into each well of 96 well plates (Falcon). All plates were air dried and one larva was added per well. Plates were sealed with pre-punched mylar, and incubated at 27°C, 70% relative humidity, and 24 hours darkness for 6 days. The number of surviving or dead insects was recorded for each concentration level at the end of the 6-day incubation period. LC_50_ values were estimated using a 3-parameter logistic model in GraphPad Prism 8.2 and used as the measurements of insecticidal activity. The threshold for the control mortality was less than or equaled to 20%. A relative activity was calculated using the formula (1)

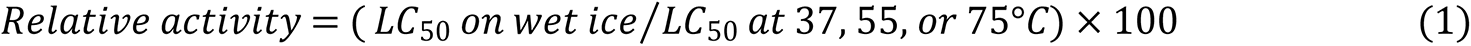

### 2.7 Acute oral toxicity assessment of Mpp75Aa1.1

Two acute oral toxicity studies were conducted in CD-1 mice, each using a study design adapted from the EPA OPPTS Guideline 870.1100. In the first study, the Mpp75Aa1.1 dose solution was formulated in a vehicle buffer (10 mM sodium carbonate/bicarbonate, pH 10.0) at a concentration of 31.7 mg Mpp75Aa1.1/ml to enable a target dose level of 2000 mg Mpp75Aa1.1/kg body weight. In addition to the test dosing solution, a protein control (bovine serum albumin, BSA) dosing solution in the same vehicle buffer was prepared at a similar concentration (31.2 mg/ml) to Mpp75Aa1.1 to enable a target dose level of 2000 mg BSA/kg body weight. In the second study, the Mpp75Aa1.1-His dose solution was formulated in a buffer (10 mM sodium carbonate/bicarbonate, pH 10.6-10.8) at a concentration of 28.8 mg Mpp75Aa1.1-His/ml to enable a target dose level of 2000 mg Mpp75Aa1.1-His/kg body weight. In addition to the test dosing solution, a protein control (bovine serum albumin, BSA) dosing solution in buffer was prepared at a similar concentration (30.6 mg/ml) to Mpp75Aa1.1-His to enable a target dose level of 2000 mg BSA/kg body weight. It is important to note, that although slightly different experimental approaches were used in these two acute toxicity studies (while both BSA and vehicle controls were used in the Mpp75Aa1.1 study, only a BSA control was used in the Mpp75Aa1.1-His study), both studies were considered suitable for hazard characterization. Prior to initiation of studies, the dosing solutions were analyzed to confirm concentration, pH, and homogeneity. Stability was confirmed by determining both the protein concentration using UV absorbance and integrity of pre- and post-dosing solutions using SDS-PAGE.

Male and female CD-1 mice were obtained from Charles Rivers Laboratories, Spencerville, Ohio and were approximately 8 weeks old with fasted body weights ranging from 27.4 to 40.4 g for males and 21.0 to 28.7 g for females at the initiation of dose administration (Day 0) for both studies. Mice were fasted approximately 3 – 4 hours prior to dose administration and between doses. Dosing solutions were administered twice (approximately 3 – 4 hours apart) over the course of a single day by oral gavage with a dose volume of 33.3 ml/kg body weight on Day 0 to groups of 10 males and 10 females and observed for 14 days thereafter. Endpoints evaluated during the dosing and observation periods included: survival, clinical observations, body weights, body weight changes, and food consumption. Following the observation period, all surviving animals were humanely euthanized by carbon dioxide inhalation on Day 14 and subjected to a complete gross necropsy. Necropsies included macroscopic examination of the carcass and musculoskeletal system; all external surfaces and orifices; cranial cavity and external surfaces of the brain; and thoracic, abdominal, and pelvic cavities with their associated organs and tissues under the supervision of a board-certified veterinary pathologist. All work was conducted in an AALAC accredited laboratory and the study protocols were reviewed by the test facility IACUC committee prior to study initiation to ensure animal welfare. Body weight, body weight gain and food consumption were statistically analyzed as described previously (15, 16).

### 2.7 Data availability

The sequences for the 5 genes and protein translations are available in Genebank. The accession number for Mpp75Aa1 nucleotide sequence is <u>MF490291.1</u> and for the protein is <u>ASY04853.1</u>.

The accession number for Mpp75Aa2 nucleotide sequence is <u>MF490290.1</u> and for the protein is <u>ASY04852.1</u>. The accession number for Mpp75Aa3 nucleotide sequence is <u>MF490289.1</u> and for the protein is <u>ASY04851.1</u>.The accession number for Mpp75Ab1 nucleotide sequence is ON007247. The accession number for Mpp75Ab2 nucleotides sequence is ON007246. The protein sequences and new nomenclature are available at the Bacterial Pesticidal Protein Resource Center (BPPRC) https://www.bpprc.org/.

## 3 RESULTS

### 3.1 Assessment of the presence of the Mpp75A protein in *B. laterosporus*

The presence of Mpp75A proteins in different *B. laterosporus* strains was assessed by immunoblot analysis (Fig. 1). The *E.coli*-produced mature form of Mpp75Aa3 (mMpp75Aa3) without the membrane transiting signal peptide was loaded in lane 2 as a control. The resulting immunoblot indicated that anti-mMpp75Aa2 antibody recognized similar immunoreactive bands migrating to the identical molecular weight position of the mMpp75Aa3 protein for proteins extracted from different strains including the BOD (Fig. 1; lane 3 & 4) and EG4227 strains (Fig. 1; lane 5 & 6).

**Figure 1.**
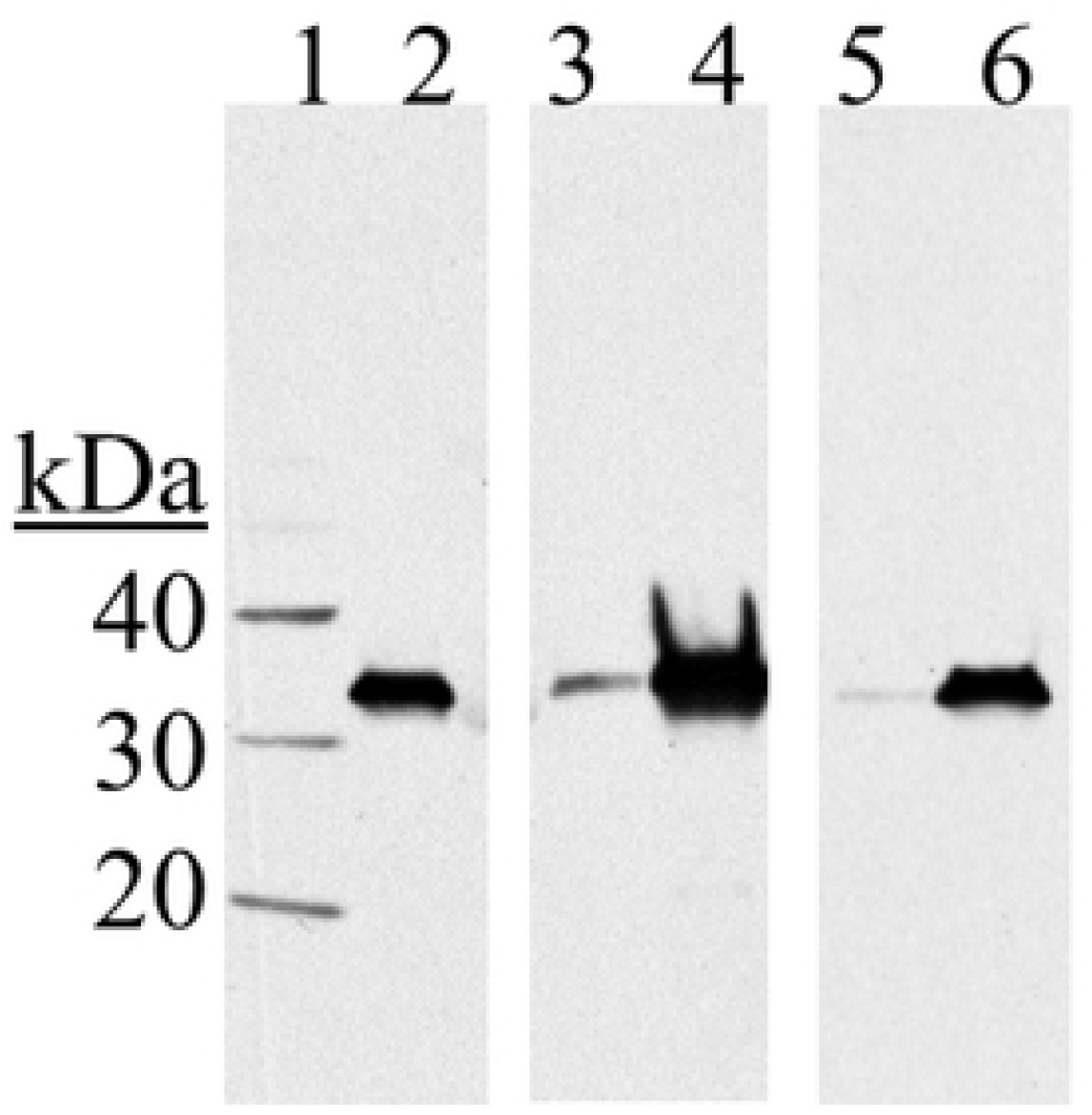
Western blot analysis of the presence of Mpp75Aa protein in different strain of *B. laterosporus*. The western blot was probed using anti-mMpp75Aa2 polyclonal antibody. Lane 1: molecular weight standard, lane 2: 15 ng of purified mMpp75Aa3 protein standard, Lane 3: whole cell extract of the BOD strain from 40 hr culture in TB, lane 4: the BOD strain cell free culture supernatant from 40 hr culture in TB, Lane 5: whole cell extract of the EG4227 strain from 40 hr culture in TB, lane 6: the EG4227 strain culture supernatant strain from 40 hr culture in TB.

**Figure 2:**
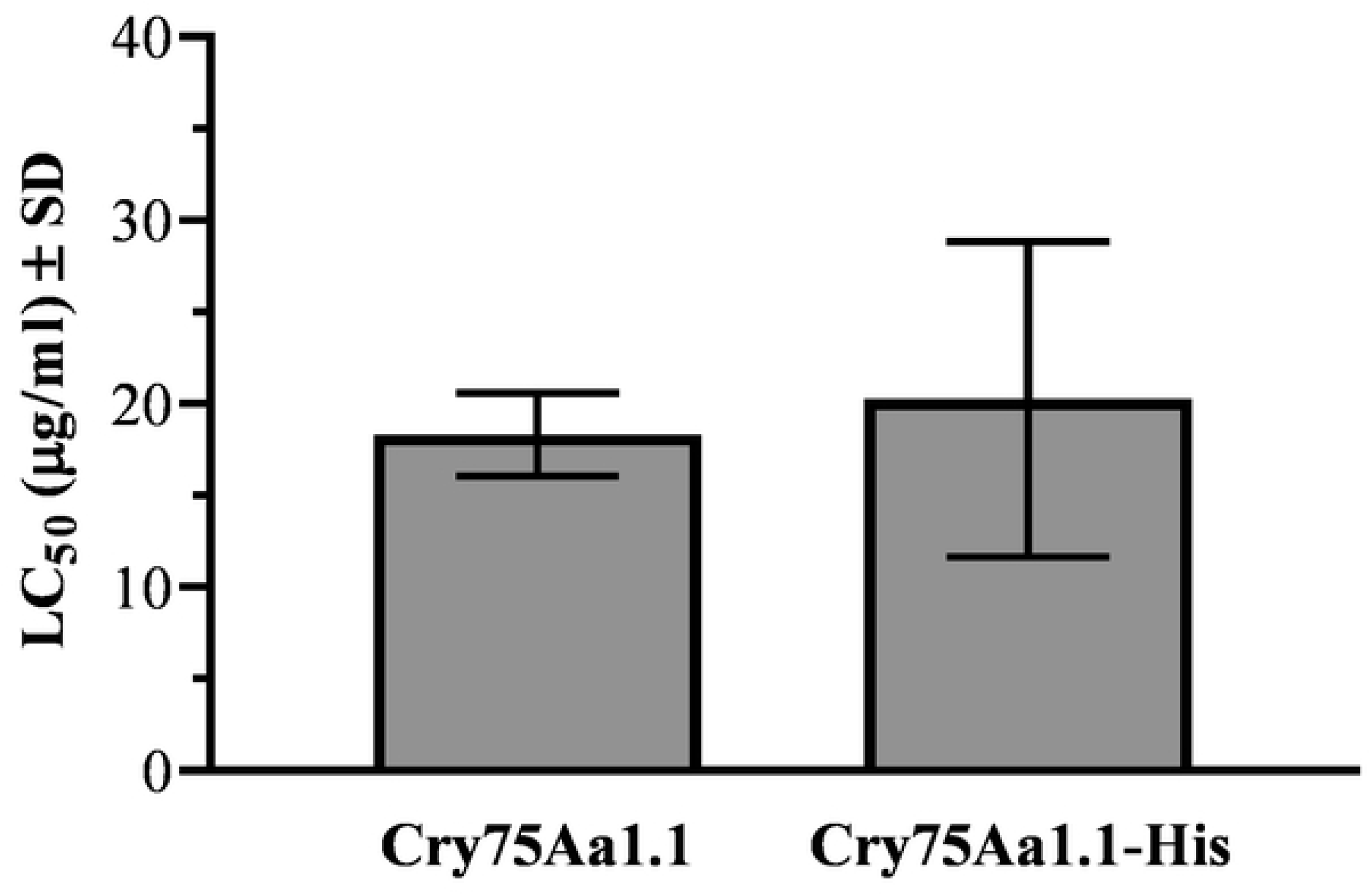
Functional activity of Mpp75Aa1.1 and Mpp75Aa1.1-His. Functional activity of protein samples was evaluated using the western corn rootworm (WCR), *Diabrotica virgifera virgifera.* All bioassays consisted of a series of 7 to 9 dilutions yielding a dose series with a two-fold separation factor ranging from 5 to 1200 μg/ml. n= 3 for Mpp75Aa1.1 and n=4 for His-tagged Mpp75Aa1.1.

No additional bands were observed in the protein samples tested, indicating that these immunoreactive bands are specific to the mMpp75Aa2 antibody. Additionally, a much stronger immunoreactive signal was detected in the cell culture medium (lanes 4 & 6) than in cell extracts (lanes 3 & 5), which is consistent with the prediction that the Mpp75Ab protein is secreted into the medium from the host cell upon the expression (11).

The Mpp75Ab gene was cloned from the BOD and EG4227strains, respectively. Two protein sequences designated as Mpp75Ab1 and Mpp75Ab2 were deduced from Mpp75Ab genes cloned. Two protein sequences were aligned with three Mpp75Aa sequences in a multiple-sequence alignment (supplemental 1) followed by a pairwise comparison (supplemental 2) and the five proteins were all at least 93% identical to each other. Therefore, both the immunoblot and protein sequence analyses confirmed that Mpp75Ab proteins exist in various strains of *B. laterosporus* including the probiotic BOD strain for human consumption.

### 3.2 Purification and Characterization of Mpp75Aa1.1

Mpp75Aa1.1 and Mpp75Aa1.1-His were purified from *E. coli* cells expressing each respective protein. Data generated from the subsequent characterization of each protein are presented in Table 1. Densitometric analysis of SDS–PAGE gels indicated that both Mpp75Aa1.1 were purified to >85% purity. Both Mpp75Aa1.1 and Mpp75Aa1.1-His displayed their expected apparent molecular weights by SDS-PAGE of ∼32 kDa and ∼33 kDa, respectively. The identities of the Mpp75Aa1.1 and Mpp75Aa1.1-His were confirmed by tryptic peptide mass fingerprinting using MS analysis, resulting in 59.5% and 68.9% coverage of the entire protein sequences, respectively. The N-terminal sequences of Mpp75Aa1.1 and Mpp75Aa1.1-His were detected, both of which are consistent with the expected sequences except for the absence of the N-terminal methionine. Removal of the N-terminal methionine by *E. coli* methionine aminopeptidase is a common modification that occurs co-translationally before completion of the nascent protein chain, and typically has no effect on protein structure or activity (45–47). Additionally, the expected C-terminal His tag (-HHHHAHHH) was identified on Mpp75Aa1.1-His.

**Table 1.**
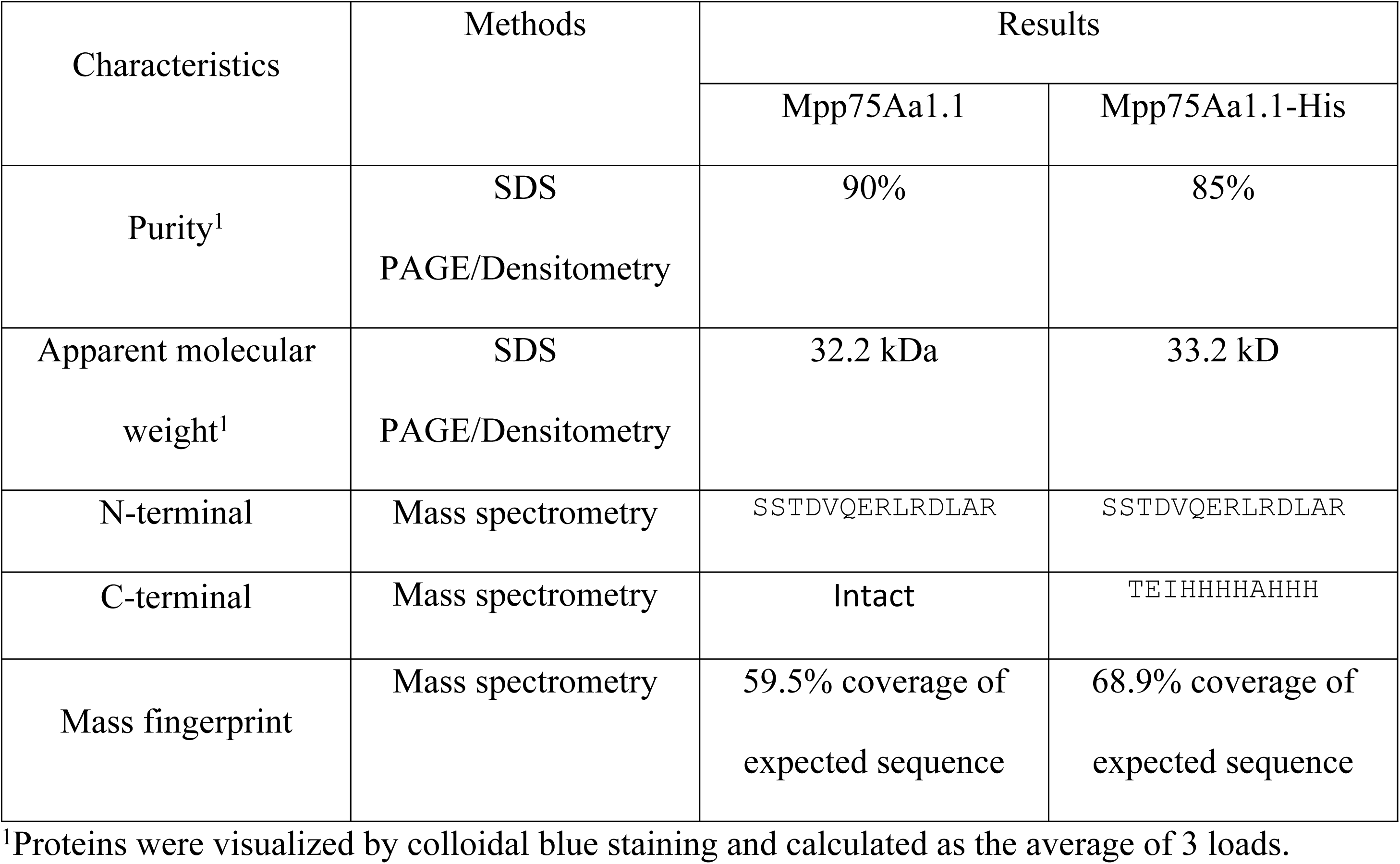
Characteristics of Mpp75Aa1.1 and Mpp75Aa1.1-His.

Both Mpp75Aa1.1 and Mpp75Aa1.1-His exhibited robust insecticidal activity against WCR (Fig 2). The LC_50_ values for Mpp75Aa1.1 and Mpp75Aa1.1-His were 18.31 ± 1.31 μg/ml and 20.25 ± 4.3 μg/ml, respectively. There was no significant difference (p=0.6556) between LC_50_ values for the Mpp75Aa1.1 and Mpp75Aa1.1-His proteins indicating that the proteins have equivalent potency and can therefore be considered equivalent, or suitable surrogates, for protein safety assessment.

### 3.3 Assessing the bioinformatic relationship of Mpp75Aa1.1 to other proteins

To evaluate the sequence similarity of Mpp75Aa1.1 and Mpp75Aa1.1-His to known proteins, a FASTA (v36.3.5d) search of all proteins in GenBank release 335 using both sequences was conducted (Table 2). This bioinformatic search resulted in identical profiles using the non-tagged or His-tagged Mpp75Aa1.1 variant as a query. Notably, the top alignment, which represents the self-identification of Mpp75Aa1.1, is labeled as “insecticidal protein” from *B. laterosporus* (NCBI Accession ASY04853.1). This annotation is unsurprising as Mpp75Aa1.1 is a known insecticidal member of the ETX_MTX2 family of proteins (11).

**Table 2.** Summary of alignments for the FASTA searches of the GenBank 335 all protein database using Mpp75Aa1.1 and Mpp75Aa1.1-His sequences.

| Sequence Name | Search of the GenBank 335 Protein Sequence Database |  |  |  |  |  |
| --- | --- | --- | --- | --- | --- | --- |
|  | # of Returned Alignments | Aligned Sequence Accession | Description | Identity | aa Overlap | E-score |
| Mpp75Aa1.1 | 535 | ASY04853.1 | insecticidal protein [Brevibacillus lateros (317 aa) | 100.0% | 294 | 1.1e-124 |
| Mpp75Aa1.1-His | 535 | ASY04853.1 | insecticidal protein [Brevibacillus lateros (317 aa) | 100.0% | 294 | 1e-124 |

### 3.4 Bioinformatics analyses of Mpp75Aa1.1 allergenicity and toxicity

Bioinformatic analyses to assess and evaluate whether or not the introduced protein shows any similarity to known toxins or allergens is seen as one of the key elements of safety evaluations for newly introduced proteins in GM crops (14, 48). The bioinformatic analyses performed on both Mpp75Aa1.1 and Mpp75Aa1.1-His demonstrated highly congruent results (Tables 3 and 4). No alignments to the COMPARE allergen database exceed the joint FAO/WHO prescribed criteria for assessing similarity to known allergens were observed for either Mpp75Aa1.1 or Mpp75Aa1.1-His. This included no 8-mer peptide matches and no sliding windows displaying 35% identity over 80 amino acids (Table 3) (12).

**Table 3.** Summary of alignments for the FASTA searches of the AD_2020 database using Mpp75Aa1.1 and Mpp75Aa1.1-His sequences.

| Sequence<br>Name | Search of the AD_2020 Sequence Database |  |  |  |  |  |  |  |
| --- | --- | --- | --- | --- | --- | --- | --- | --- |
|  | FASTA search |  |  |  |  |  |  |  |
|  | 8 mer<br>alignmen<br>t | 35%<br>ID 80<br>aa | # of<br>Returned<br>Alignments | Aligned<br>Sequence<br>Accession | Description | Identity | aa<br>Overlap | *E*-<br>score |
| Mpp75Aa1.1 | No | No | 7 | ACZ57582.1 | Putative Ole e 11.0101 allergen precursor (364 aa) | 20.2% | 272 | 0.17 |
| Mpp75Aa1.1-His | No | No | 7 | AGS80276.1 | Putative manganese superoxide dismutase [A (191 aa) | 21.0% | 119 | 0.089 |

Searches against the toxin database (TOX_2020) resulted in the same observation for each of the query sequences (Table 4), with each of them displaying an *E-*score of ≤ 1e-5 (1 x 10^-5^), the cutoff which has been used as a conservative threshold for alignment significance (38, 39, 43). The observed alignment to the Mpp75Aa1.1 variant displayed 25.9% sequence identity to “Epsilon-toxin type B” from *C. perfringens* (Q02307).

**Table 4.** Summary of alignments for the FASTA searches of the TOX_2020 database using Mpp75Aa1.1 and Mpp75Aa1.1-His sequences.

| Sequence<br>Name | Search of the TOX_2020 Sequence Database |  |  |  |  |  |
| --- | --- | --- | --- | --- | --- | --- |
|  | # of Returned<br>Alignments | Aligned Sequence<br>Accession | Description | Identit<br>y | aa<br>Overlap | *E*-score |
| Mpp75Aa1.1 | 2 | sp\|Q02307\|ETXB_CL<br>OPF | Epsilon-toxin type B OS=Clost | 25.9% | 297 | 9.6e-23 |
| Mpp75Aa1.1-His | 4 | sp\|Q02307\|ETXB_CL<br>OPF | Epsilon-toxin type B OS=Clost | 25.9% | 297 | 7.8e-23 |

### 3.5 Assessment of the susceptibility of the Mpp75Aa1.1 to degradation by pepsin

Degradation of Mpp75Aa1.1 by pepsin was assessed using an established and standardized assay (49). The resulting SDS-PAGE gels indicated that the intact Mpp75Aa1.1 (Fig. 3A) or Mpp75Aa1.1-His (Fig. 3B) protein was completely disappeared after incubation with pepsin at 37 °C for 0.5 min (Fig. 3), the first time-point tested. The limit of detection (LOD) of Mpp75Aa1.1 by staining with Colloidal Brilliant Blue G was observed at approximately 5 ng or approximately 0.5% of total Mpp75Aa1.1 or Mpp75Aa1.1-His loaded. This demonstrates that at least 99.5% of Mpp75Aa1.1 or Mpp75Aa1.1-His was degraded by pepsin within 0.5 min. Additionally, for both forms of Mpp75Aa1.1, a smaller Mpp75Aa1.1 peptide fragment of ∼4 kDa was observed at the 0.5 min time point but was completely degraded after 5 min of incubation with pepsin at 37 °C (Fig. 3, Lanes 5-8). There was no change in the protein banding pattern for the protein when incubated at 37 °C in the absence of pepsin, indicating that the observed protein degradation was a direct result of the proteolytic activity of pepsin and not due to instability of the protein when incubated at 37 °C (Fig. 3, Lanes 3 and 12). Additionally, there was no change in the protein band corresponding to pepsin (∼38 kDa) when incubated at 37 °C in the absence of Mpp75Aa1.1 or Mpp75Aa1.1-His indicating that pepsin was stable at 37 °C over the course of the experiment (Fig. 3, Lanes 2 and 13).

**Figure 3:**
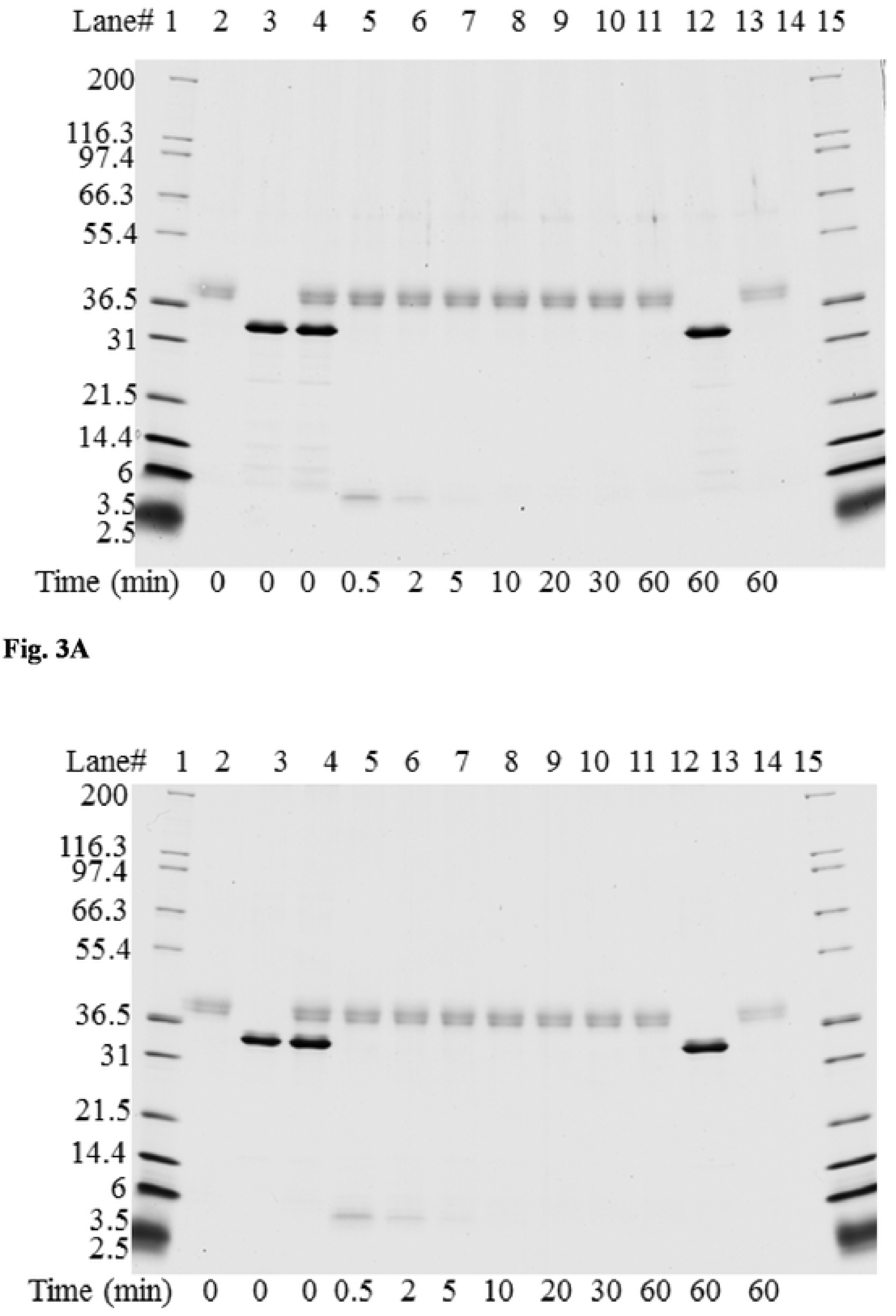
SDS-PAGE analysis of degradation of Mpp75Aa1.1 and Mpp75Aa1.1-His by pepsin. Degradation of Mpp75Aa1.1 and His-tagged Mpp75Aa1.1 upon exposure to pepsin was assessed by analysis of SDS-PAGE gels after staining with Colloidal Brilliant Blue G protein stain. Based on the pre-reaction protein concentration of Mpp75Aa1.1 and His-tagged Mpp75Aa1.1, 1 µg of was loaded in each lane containing Mpp75Aa1.1 (Fig. 3A) or His-tagged Mpp75Aa1.1 (Fig. 3B) (Lanes 3-12). After the addition of pepsin at an approximate ratio 1 μg total protein:10 U of pepsin (Lane 4) and incubation at 37°C (Lanes 5-11), the full length Mpp75Aa1.1 or His-tagged Mpp75Aa1.1 is completely degraded. Mpp75Aa1.1, His-tagged Mpp75Aa1.1 or pepsin incubated independently at 37°C for the duration of the experiment (Lanes 12 and 13, respectively) exhibited no change in band intensity. Approximate molecular weights (kDa) are shown on the left side of the gel and correspond to the molecular weight markers loaded.

### 3.6 Heat lability of the Mpp75Aa1.1

The thermal stability of Mpp75Aa1.1 and Mpp75Aa1.1-His was determined by comparing LC_50_ values of WCR in diet overlay bioassays when Mpp75Aa1.1 and Mpp75Aa1.1 were incubated at 37, 55, or 75°C ± 2 for 15 min (Fig. 4). The 37°C heat treatment did not significantly change activity compared with samples on wet ice for both Mpp75Aa1.1 (p=0.2633) and Mpp75Aa1.1-His (p=0.3659). However, when heated to temperatures of 55 and 75°C for 15 min, both Mpp75Aa1.1 and Mpp75Aa1.1-His lost at least 96.1% activity. The results confirm that Mpp75Aa1.1-His had the same pattern of heat lability as the Mpp75Aa1.1 (p=0.6556).

**Figure 4:**
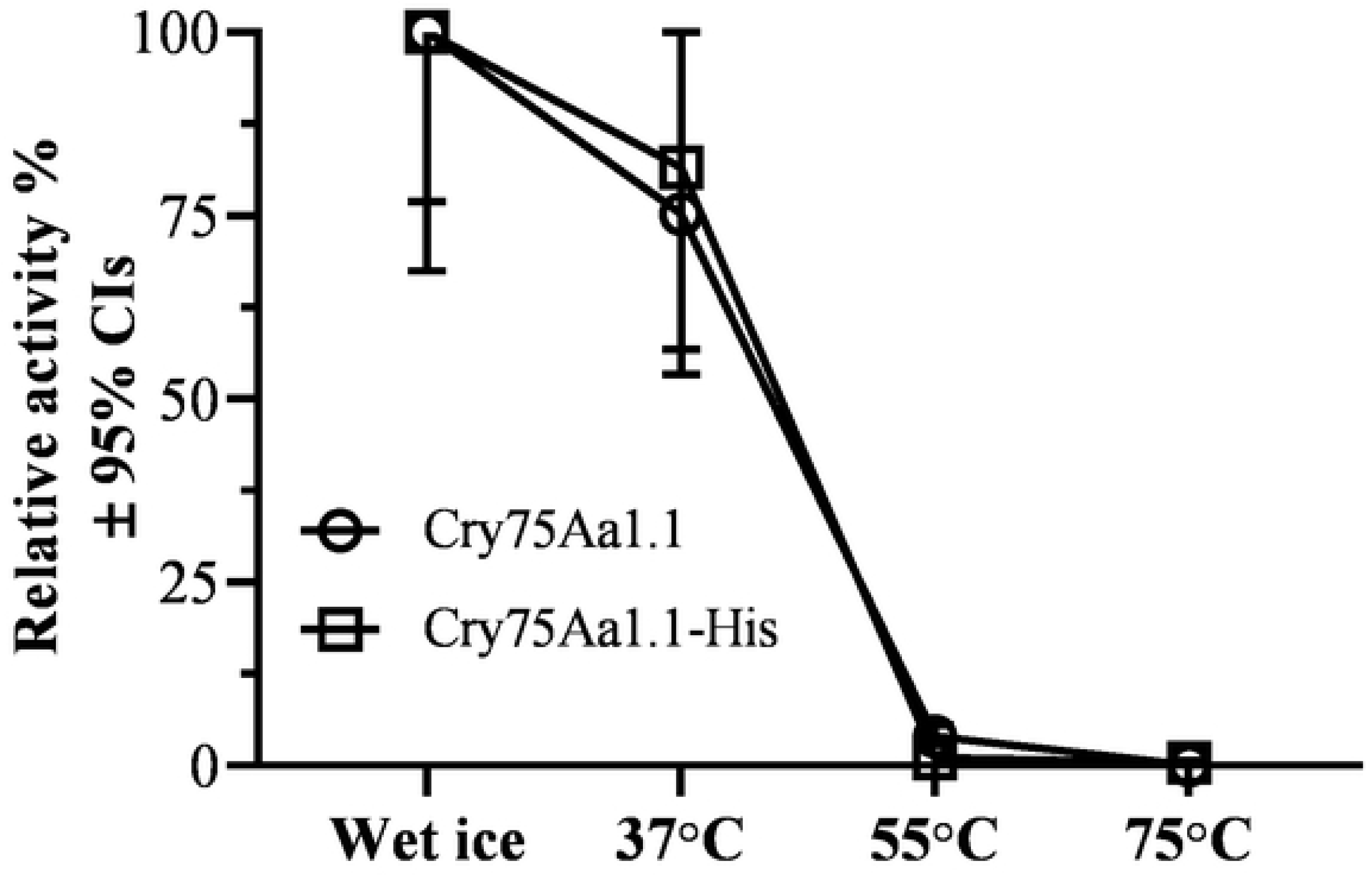
Heat lability evaluation of Mpp75Aa1.1 and Mpp75Aa1.1-His by functional assay. Relative activity of Mpp75Aa1.1 (open circle) and Mpp75Aa1.1-His (open square) after heat treated at 37, 55, and 75°C for 15 min was calculated by the LC_50_ values for the treatment on wet ice divided by the LC_50_ values for the treatment at 37, 55 and 75°C. LC_50_ values were estimated from dose-response curves of Mpp75Aa1.1 measured in 6-day diet-overlay bioassays using the western corn rootworm (WCR), *Diabrotica virgifera virgifera.* The overlapped relative activity curves illustrate there was no significant difference in heat lability between Mpp75Aa1.1 and Mpp75Aa1.1-His (p=0.6556).

### 3.7 Assessment of oral toxicity of Mpp75Aa1.1

For both Mpp75Aa1.1 and Mpp75Aa1.1-His, Tier I assessments (bioinformatics) illustrated above indicated a lack of apparent hazards and thus a higher tier assessment such as acute toxicity testing was not scientifically warranted. Furthermore, as described above, the His tag did not impact Mpp75Aa1.1 digestibility, heat stability or functional activity suggesting that Mpp75Aa1.1-His was functionally equivalent to Mpp75Aa1.1. Nevertheless, acute oral toxicity studies in mice were conducted on Mpp75Aa1.1 and Mpp75Aa1.1-His in order to provide further confirmation of mammalian safety and to determine if there were changes in the safety assessment as a result of the addition of the His tag. Mice in test groups were dosed with either Mpp75Aa1.1 or Mpp75Aa1.1-His at actual doses of 2120 and 1920 mg/kg, respectively. Following dosing, mice were observed for 14 additional days, humanely euthanized, and subsequently subjected to a macroscopic examination of their anatomy. No mortality occurred during either of these studies and no test substance-related clinical signs were observed during the studies. There were no test substance-related differences in mean body weights or mean body weight gains in either of these studies (Tables 5 & 6). There were no statistically significant differences in these endpoints for Mpp75Aa1.1-His (Tables 5 & 6). There was a significantly higher mean body weight for the test group females compared to the vehicle control females on Study Days 0 and 7 for Mpp75Aa1.1 (Table 5); however, this difference was considered incidental as the body weight difference was small, there was no difference in body weights at Day 14, there were no differences relative to the BSA control at any time point, and these differences were only observed in females. Furthermore, there were no statistically significant differences in body weight gains in this study (Table 6) and overall, an increase in body weight does not suggest toxicity. There were no test substance-related differences in food consumption observed in either of these studies (Table 7). There were no statistically significant differences in this endpoint for Mpp75Aa1.1-His (Table 7). ForMpp75Aa1.1, there was a statistically significant increase in food consumption for the test group females compared to the BSA control females for Study Days 0-7 (Table 7). This increase in food consumption was considered incidental as there were no differences relative to vehicle control, the difference was not observed from Study Days 7-14, and this difference was only observed in females. No test substance-related gross pathology findings were observed at necropsy. These results contribute to the weight of evidence and further support the conclusion that both Mpp75Aa1.1 and Mpp75Aa1.1-His and present no hazard when consumed in food or feed.

**Table 5.** Summary of acute toxicity study body weight results for both Mpp75Aa1.1 and Mpp75Aa1.1-His.

| Group | | Mean BW<br>Day 0 (grams) ± SD | | | Mean BW<br>Day 7 (grams) ± SD | | | Mean BW<br>Day 14 (grams) ± SD | | |
| --- | --- | --- | --- | --- | --- | --- | --- | --- | --- | --- |
|  |  | Test | Vehicle | BSA | Test | Vehicle | BSA | Test | Vehicle | BSA |
| Mpp75Aa1.1 | Males | 35.5 ± 1.9 | 35.4 ± 2.0 | 33.8 ± 3.3 | 37.5 ± 2.3 | 36.3 ± 3.7 | 35.6 ± 3.2 | 38.5 ± 2.6 | 37.5 ± 3.1 | 36.3 ± 3.1 |
| | Females | 27.1 ± 0.7<sup>a</sup><br>† | 26.1 ± 1.0 | 27.0 ± 1.0 | 29.3 ± 0.8<sup>b</sup><br>† | 27.9 ± 1.5 | 28.3 ± 1.8 | 30.7 ± 0.9 | 29.5 ± 2.2 | 30.0 ± 1.6 |
| Mpp75Aa1.1<br>-His | Males | 30.9 ± 2.2 | NA | 30.8 ± 1.3 | 32.5 ± 2.4 | NA | 33.0 ± 1.6 | 33.0 ± 2.7 | NA | 33.4 ± 2.0 |
| | Females | 23.6 ± 0.9 | NA | 23.9 ± 1.6 | 25.3 ± 1.2 | NA | 26.0 ± 1.6 | 26.1 ± 2.1 | NA | 26.8 ± 2.1 |

<sup>a</sup> = Statistically significant difference by ANOVA with Dunnett's Test at $p < 0.05$ .
<sup>b</sup> = Statistically significant difference by Kruskal-Wallis Test with Dunn's Test at $p < 0.05$ .
† = Statistical comparison between test and vehicle.
BW = body weight; SD = Standard Deviation; BSA = Bovine Serum Albumin protein control.
N = 10 animals/sex/dose.
NA = not applicable as this treatment group was not present in study.
Dose levels for Mpp75Aa1.1 and Mpp75Aa1.1-His were 2120 and 1920 mg protein/kg body weight, respectively.

**Table 6.** Summary of acute toxicity study body weight changes for both Mpp75Aa1.1 and Mpp75Aa1.1-His.

| Group | | Mean BW Change<br>Days 0-7 (grams) ± SD | | | Mean BW Change<br>Days 7-14 (grams) ± SD | | | Mean BW Change<br>Days 0-14 (grams) ± SD | | |
| --- | --- | --- | --- | --- | --- | --- | --- | --- | --- | --- |
|  |  | Test | Vehicle | BSA | Test | Vehicle | BSA | Test | Vehicle | BSA |
| Mpp75Aa1<br>.1 | Males | 2.0 ± 1.1 | 0.9 ± 2.2 | 1.8 ± 0.9 | 1.0 ± 0.5 | 1.2 ± 1.3 | 0.8 ± 0.4 | 3.0 ± 1.5 | 2.1 ± 1.8 | 2.6 ± 0.7 |
| | Females | 2.2 ± 1.0 | 1.8 ± 0.7 | 1.3 ± 1.2 | 1.4 ± 1.1 | 1.7 ± 1.0 | 1.7 ± 1.0 | 3.7 ± 1.2 | 3.4 ± 1.4 | 3.0 ± 1.2 |
| Mpp75Aa1<br>.1-His | Males | 0.3 ± 0.7 | NA | 0.8 ± 0.9 | NA | NA | NA | 0.8 ± 1.0 | NA | 1.2 ± 1.3 |
| | Females | 0.6 ± 0.8 | NA | 1.0 ± 1.5 | NA | NA | NA | 1.5 ± 1.6 | NA | 1.8 ± 1.6 |

No statistically significant differences were observed by ANOVA with Dunnett's Test at $p < 0.05$ .
BW = body weight; SD = Standard Deviation; BSA = Bovine Serum Albumin protein control.
N = 10 animals/sex/dose.
NA = not applicable as this treatment group was not present in study.
Dose levels for Mpp75Aa1.1 and Mpp75Aa1.1-His were 2120 and 1920 mg protein/kg body weight, respectively

**Table 7.** Summary of acute toxicity study food consumption results for both Mpp75Aa1.1 and Mpp75Aa1.1-His.

| Group |  |  | Mean Food Consumption<br>Days 0-7 (grams/animal/day) ± SD |  |  | Mean Food Consumption<br>Days 7-14 (grams/animal/day) ± SD |  |  |
| --- | --- | --- | --- | --- | --- | --- | --- | --- |
|  |  |  | Test | Vehicle | BSA | Test | Vehicle | BSA |
| Mpp75Aa1.1 | Males | Mean ± SD | 6.0 ± 1.2 | 5.2 ± 1.0 | 6.5 ± 2.6 | 5.2 ± 0.7 | 5.1 ± 0.6 | 5.2 ± 0.9 |
|  |  | N | 10 | 10 | 10 | 10 | 10 | 10 |
|  | Females | Mean ± SD | 7.4 ± 2.9<sup>b, #</sup> | 7.3 ± 3.1 | 5.0 ± 1.8 | 6.8 ± 2.5 | 6.2 ± 2.5 | 5.3 ± 2.1 |
|  |  | N | 10 | 10 | 10 | 9 | 9 | 10 |
| Mpp75Aa1.1-HIS | Males | Mean ± SD | 4.9 ± 1.2 | NA | 5.5 ± 0.9 | 5.7 ± 1.8 | NA | 5.5 ± 1.2 |
|  |  | N | 10 | NA | 10 | 10 | NA | 10 |
|  | Females | Mean ± SD | 5.2 ± 1.5 | NA | 5.2 ± 1.5 | 5.5 ± 1.3 | NA | 4.9 ± 1.4 |
|  |  | N | 9 | NA | 9 | 8 | NA | 9 |

<sup>b</sup> = Statistically significant difference by Kruskal-Wallis Test with Dunn's Test at p < 0.05.
<sup>#</sup> = Statistical comparison between test and BSA.
BW = body weight; SD = Standard Deviation; BSA = Bovine Serum Albumin protein control.
NA = not applicable as this treatment group was not present in study.
N = 10, some cage data not collected due to excessive food spillage.
Dose levels for Mpp75Aa1.1 and Mpp75Aa1.1-His were 2120 and 1920 mg protein/kg body weight, respectively.

## 4 DISCUSSION

The focus of the present study was to conduct a comprehensive assessment of the food and feed safety for Mpp75Aa1.1 using the tiered weight of evidence approach. In addition, this study also evaluated the feasibility using a His-tagged protein for protein safety assessment (13, 48). Prior to any experimental assessment of the safety of a protein in food or feed, an evaluation of HOSU for both the source organism (i.e., the organism from which the protein is derived) and the protein itself can be determined (13). The Mpp75Aa1.1 protein was derived from *B. laterosporus*, an endospore-forming insecticidal *bacillus* not commonly associated with human disease (50–52). *B. laterosporus* is an abundant organism that has been isolated from a wide range of environments including soil, rocks, dust, and both fresh and sea waters (53–55). Furthermore, *B. laterosporus* is found in many foods such as cheese (56), curd (55), beans (57), and honey (58) as well as being ingested in some commercial probiotics (59) and feed additives (60).

Morphological examination, genome sequencing and analysis of the 16S rRNA identifies that the source organisms for Mpp75Aa1 (EG5553) and Mpp75Aa3 (EG5552) are phylogenetically related to the *B. laterosporus* BOD strain that is commercially available as Latero-Flora^TM^ (Global Healing Institute, Houston, TX) human dietary supplement (11). The presence of the Mpp75Ab1 protein in human dietary supplements was demonstrated using immunoblotting analysis using the Mpp75Aa2 antibody as well as the amino acid sequence alignment (Fig. 1 & supplemental data), indicating that humans have been directly exposed to the Mpp75Ab protein through the use of dietary supplements. Taken together, the widespread presence of *B. laterosporus* in the environment, and the presence of the Mpp75A protein in food and feed, provide support for a history of safe consumption by humans and other vertebrate animals.

Bioinformatic analyses identifies the Mpp75Aa1.1 protein as an insecticidal member of the ETX_MTX2 β-PFP family, which has been described previously (18). The presence of ETX_MTX2 proteins in foods has also been confirmed in various vegetables and food crops with a history of safe consumption, such as spinach (*Spinacia oleracea*), sugar beet (*Betula vulgaris*), wheat (*Triticum aestivum*), cucumber (*Cucumic sativus*) and blue catfish (*Ictalurus furcatus*) (19). In addition, the safety of β-PFPs in crops used for food and feed is supported by the safe consumption of the binary toxin complex of Tpp35Ab1 (20, 21), and its partner protein Gpp34Ab1 (19). The recent safety assessment of three Mpp51Aa2 protein variants provides further verification for the safe consumption of ETX_MTX2 β-PFP proteins (15). Furthermore, ETX_MTX2 proteins in biopesticide products have a global HOSU in the control of mosquitoes and black flies (19, 61).

In addition to providing information regarding protein identity, bioinformatic analysis can also be used to assess potential allergenicity or toxicity of a protein by demonstrating whether a protein shares primary sequence homology, and therefore potential higher order structural homology, to known allergens or toxins. Results from these analyses demonstrate no relevant similarity between Mpp75Aa1.1 and any known allergenic protein in the COMPARE database (Table 3), and single significance threshold exceeding alignments in TOX_2020 (Table 4). As noted previously, Mpp75Aa1.1 is a member of the ETX_MTX2 β-PFP family and therefore shares a similar overall structure with other ETX_MTX2 family members, including some mammalian toxins such as epsilon toxin and aerolysin (Table 4). Although ETX_MTX2 β-PFPs share common structural characteristics with epsilon toxin, divergence in key amino acid sequences and functional domain structures provides discreet organism specificity among the protein family members. Previous analyses of this protein class have demonstrated that ETX_MTX2 β-PFPs consist of three main structural domains that are generally conserved amongst family members even if the level of primary sequence identity between members can be as low as 25% (19, 62). One domain is responsible for receptor-binding/target-specificity while the other two domains contain the pore-forming loop and oligomerization regions responsible for the beta-pore forming function. More important, the pore-formation and oligomerization domains share a higher degree of structural conservation within the family (19, 22–25, 62, 63). The increased divergence in amino acid sequence of the receptor binding domain relative to the rest of the protein underscores its importance in providing specificity to β-PFPs, which has been illustrated by amino acid substitutions (64–66). Replacement of two surface exposed tyrosine residues in the head region of epsilon toxin, reduced cell binding and cytotoxic activities in MDCK.2 cells and abolished toxicity in mice (67), indicating that epsilon toxin specificity is conveyed by the surface-exposed amino acid residues within the receptor binding domain. Such domain-based protein characterizations reveal that ETX_MTX2 proteins have distinct activation processes despite their overall structural similarity (19). The above data clearly indicate that identifying homologous ETX_MTX2 family members in TOX_2020 database warrants further investigation but does not by itself demonstrate toxicity. This highlights the importance of incorporating a domain-based evaluation into protein safety assessments when using the weight of evidence approach (19).

Although an evaluation of HOSU and bioinformatic analyses provide evidence that Mpp75Aa1.1 is safe for consumption by humans or other vertebrate animals, the weight of evidence used in assessing the safety of a protein was also evaluated through empirical experimentation. Evaluation of the susceptibility of the protein of interest to enzymatic digestion is an important element of the safety assessment for introduced proteins as most dietary proteins are digested to constituent amino acids and small peptides in the gastrointestinal system and absorbed for nutritive purposes. Rapid degradation by gastric enzymes provides evidence that exposure to the intact introduced protein will be minimized following consumption and is therefore less likely to be allergenic (12, 14, 48). When empirically tested, the intact Mpp75Aa1.1 and His-tagged Mpp75Aa1.1 were both rapidly degraded by pepsin (not detectable after 0.5 min incubation), and a small (∼4 kDa) transiently-stable fragment of the protein was completely disappeared after 5 min by pepsin (68) which is consistent with the susceptibility to pepsin by other ETX_MTX2 proteins (15). Thus, Mpp75Aa1.1 is susceptible to mammalian digestive enzymes, indicating that exposure to structurally and functionally intact Mpp75Aa1.1 is unlikely when consumed as part of food or feed. The presence of a His tag on the C-terminal end of the protein had no impact on the susceptibility of Mpp75Aa1.1 to pepsin suggesting that the use of His-tagged proteins is appropriate for use in digestion studies

The Mpp75Aa1.1 showed a complete loss of functional activity after heating at temperatures at or above 55°C for 15 min. These observations were consistent with the extensive testing of other Cry proteins in response to heating (5, 15, 69, 70) Given virtually all consumed foods from maize are exposed to heating during processing or cooking for humans (71), the heat lability of Mpp75Aa1.1 is consistent with the conclusion that dietary exposure to functionally intact Mpp75Aa1.1 is unlikely. Mpp75Aa1.1 heat lability results and rapid degradation by digestive enzymes are consistent with the previous studies conducted for a different member of the ETX_MTX2 family (15). The presence of a His tag on the C-terminal end of Mpp75Aa1.1 had no impact on the heat lability of Mpp75Aa1.1, equally suggesting that the use of His-tagged proteins is appropriate in heat lability studies The weight of evidence from the first tier assessment leveraging the domain-based characterization, and the demonstration that there is negligible potential for exposure to the intact protein supports that the Mpp75Aa1.1, and Mpp75Aa1.1-His-tagged proteins are not a hazard to humans or other vertebrate animals (14). Nevertheless, evaluation of the potential for toxicity of Mpp75Aa1.1 was still examined with the second tier approach. Acute toxicity studies were conducted in mice with both forms of Mpp75Aa1.1 to confirm a lack of toxicity through acute mechanisms (72, 73). Protein acute toxicity tests, with doses of 1920 mg Mpp75Aa1.1-His/kg body weight and 2120 mg Mpp75Aa1.1/kg body weight, resulted in no impacts on mice, which is consistent with the extensive testing of other ETX_MTX2 proteins or other β−PFPs showing no evidence of toxicity towards humans or other vertebrate animals even though they share overall structural similarity to mammalian toxins (5, 15, 21). The presence of a His tag had no impact on the mammalian safety of Mpp75Aa1.1 suggesting that the use of His-tagged proteins is appropriate for use in mouse acute toxicity studies. Thus, the toxicological study results are consistent with the conclusions of safety based upon functional domain-based characterization (18, 19).

One of the most difficult challenges for conducting the safety assessment of proteins is producing a sufficient amount of high-purity protein to conduct assessments. The use of affinity tags such as His tags can simplify protein purification, especially for intractable or difficult to express/purify proteins (27, 74). Until now, the impact of a His tag on protein safety assessment has not been evaluated explicitly. In this study, Mpp75Aa1.1 with the C-terminal His tag was produced from *E. coli*. and purified using the His tag affinity technology. Results from analyses of functional activity, digestibility, heat stability and acute toxicity demonstrate that the His-tag did not impact the Mpp75Aa1.1 safety profile and therefore the use of a His-tag protein for Mpp75Aa1.1 can be considered as suitable for safety assessment. More broadly, when equivalent physiochemical and functional properties between a protein and its affinity-tagged variant are established the use of such affinity-tagged variants should be suitable for assessing the safety of the protein of interest.

In summary, the Mpp75Aa1.1 protein was produced, characterized, and assessed for safety. There was no indication of a hazard to humans and other vertebrate animals consuming foods containing Mpp75Aa1.1 at a dose up to 2120 mg/kg body weight, consistent with the domain-based characterization. Furthermore, the presence of a His tag did not impact the safety assessment of Mpp75Aa1.1 suggesting that future protein safety studies can be conducted using His-tagged proteins.

## ACKNOWLEDGMENTS

The authors would like to thank Dr. John Vicini and Graham Head for critical reading of the manuscript.

